# Single cell tracking based on Voronoi partition via stable matching

**DOI:** 10.1101/2020.08.20.259408

**Authors:** Young Hwan Chang, Jeremy Linsley, Josh Lamstein, Jaslin Kalra, Irina Epstein, Mariya Barch, Kenneth Daily, Phil Synder, Larsson Omberg, Laura Heiser, Steve Finkbeiner

## Abstract

Live-cell imaging is an important technique to study cell migration and proliferation as well as image-based profiling of drug perturbations over time. To gain biological insights from live-cell imaging data, it is necessary to identify individual cells, follow them over time and extract quantitative information. However, since often biological experiment does not allow the high temporal resolution to reduce excessive levels of illumination or minimize unnecessary oversampling to monitor long-term dynamics, it is still a challenging task to obtain good tracking results with coarsely sampled imaging data. To address this problem, we consider cell tracking problem as “stable matching problem” and propose a robust tracking method based on Voronoi partition which adapts parameters that need to be set according to the spatio-temporal characteristics of live cell imaging data such as cell population and migration. We demonstrate the performance improvement provided by the proposed method using numerical simulations and compare its performance with proximity-based tracking and nearest neighbor-based tracking.

## I. Introduction

The time-lapse imaging technique is a powerful method used to record the molecular/functional dynamics of individual cells and their migration pattern. To understand these processes of an individual cell, it is required to detect cells and link individual tracks over time. There exist several object tracking tools including [1]–[7] and they can be broadly categorized into two groups: i) tracking-by-contour-evolution methods and ii) tracking-by-detection methods [8]. Tracking-by-contour-evolution methods [6], [7], [9]–[11] solve the segmentation and tracking tasks simultaneously by segmenting the objects in the earlier frame and evolving their contour in consecutive frames. Tracking-by-detection methods first segment the cells in all frames and later link temporal associations between the segmented object using mostly probabilistic models. Since this approach decouples the detection and the tracking processes, it is computationally efficient and thus widely used in multi-cell tracking [12].

A scenario of high temporal resolution and/or low cell migration speed (or small movement between consecutive frames) allows relative simple identification of the correspondence between cells for tracking. For example, one can start by segmenting the cells in the first frame of a video and evolve their contours in consecutive frames [13], [14]. However, often biological experiment does not allow the high temporal resolution to minimize unnecessarily oversampling time point or exposing the cells to the excessive level of illumination to measure these processes of individual cells over days [15]. Thus, a scenario of low temporal resolution and/or high cell speed (or large movement) is a more practical and challenging problem since this setting complicates the identification of the correspondence between cells. Since existing tracking methods with the low temporal resolution are not robust to cell movements, it often requires significant manual interventions or semi-automated solution [5].

In the field of computer vision, decades of research on visual object tracking have developed a diverse set of approaches and provided good solutions but tracking generic objects has remained challenging [16]. As cells generally often show deformation over time or they have similar morphology, traditional visual tracking methods are not suitable for tracking the non-rigid object. To handle these issues, we need to overcome numerous inevitable factors such as shape and scale change, variation and pose change, partial occlusion, and illumination effect. Recently, deep learning methods demonstrate powerfully extract multi-level feature representations from the raw image and provide good performance for both visual tracking and saliency detection [17]–[20]. However, these deep learning approaches often require a large training dataset and it is challenging to acquire a large manually curated tracking dataset from live-cell imaging due to the heterogeneity of cell morphology and different behavior across different experiments and perturbations.

Here, we focus on tracking-by-detection method, especially distance-based tracking method among a diverse set of different approaches of object tracking. First, we introduce Voronoi partition, a geometric naturalistic method to determine neighbors in a set of objects [21], and use it as a robust and reliable metric to identify a mappable condition, instead of using the overlap of objects in consecutive frames or nearest distance metric. Second, motivated by a stable matching problem [22], we consider tracking process as a matching problem to link individual tracks over time. We show that Voronoi-based preference metric enables stable matching by removing *blocking object* which will be defined in the paper.

## II. Background

In general, tracking-by-detection methods comprise two parts: i) cell detection or segmentation and ii) linking the objects. For cell detection, the methods range from simple intensity-based thresholding, edge detection or shape matching to a more sophisticated algorithm such as active contour, morphological operation, machine learning and deep learning applications [8], [23]. Cell tracking methods range from a simple overlap-based label propagation, contour evolution, distance-based or nearest neighbor-based linking, graph-based approach to probability-based or multiple hypothesis-based approach [8]. Since tracking-by-detection method comprises two parts, i.e., the detection and the tracking processes, both detection error and linking error will cause the overall object tracking errors. However, since recent deep learning approaches demonstrate robust segmentation performance in many biomedical imaging applications, here, we mainly focus on the tracking process.

### A. Related work: distance-based tracking approaches

If the cell movement is small in consecutive frames, we can simply check the overlap region of segmented cell masks in consecutive frames. Similarly, one can use the proximity distance or the nearest neighbors in space and time. we can consider that two cells are mappable to each other.

Overlap-based tracking encodes each cell as the same object at the next time point based on the overlap of the coordinates. Cell overlap between consecutive frames is important for correctly tracking the cells, as many algorithms rely on this overlap [8]. Proximity-based tracking encodes an object as the same object at the next time point based on the proximity of the coordinates of the centroid of its segmented object mask to a segmented object mask at the previous time point. Nearest neighbor-based tracking or adjacency-based tracking is a variant on the proximity-based tracking where it encodes an object as the same object at the next time point based on the nearest object to a segmented object mask at the previous time point instead of using the (predefined) proximity radius.

One can use neighbor information of individual cell as a graph (i.e., each vertex in graph represents an individual cell and an edge represents distance-based nearest neighbor) and if the neighbor information does not change much over time, then one can map to each other based on this information. However, if cells undergo significant movement or cells are located in dense population region or neighbor information changes between consecutive frames (i.e., cell migration, differentiation or death), we expect that cell tracking methods based on the neighbor information perform poorly.

### B. Stable marriage problem

We will consider the cell tracking problem as a stable matching problem to assign individual objects for tracking similar to that of the Gale-Shapley algorithm for the stable marriage problem [24], [25]. The stable marriage problem (also known as a stable matching problem) is the problem of finding a stable matching between two equally sized sets of elements given an ordering of preferences for each element, for instance, the best-known example is the assignment of graduating medical students to their first hospital appointments [26]. This is distinct from the stable roommates problem [27] which allows matches between any two elements but in the stable marriage problem, there exist two classes that need to be paired with each other. For instance, in the cell tracking process, we could consider objects in the consecutive time frame as two classes respectively.

In this setting, a matching is a bijection from the elements (*a*_*i*_) of one set (*a*_*i*_ ∈ *A*) to the elements (*b*_*r*_) of the other set (*b*_*r*_ ∈ *B*) and a matching is stable when there does not exist any match (*A, B*) which both prefer each other to their current partner under the matching [28]. As an example, a matching is *not stable* if:

- *∃a*_*i*_ where the first matched set prefers some given element *b*_*r*_ of the second matched set over the element to which *a*_*i*_ is already matched, and
- *b*_*s*_ also prefers *a*_*k*_ over the element to which *b*_*t*_ is already matched.

Here, we consider objects in the frame at *t* and *t* + 1 as two classes (*A, B*) in *stable matching* problem and the difference is that, in our setting, two classes may have different number of elements (i.e., we may have *n* objects for *A* and *m* objects for *B* respectively). Therefore, the cell tracking process concerns a more general variant of the stable matching problem differing in the following ways from the basic *n*−to−*n* form of the stable marriage problem:

- each object at time step *t* may only be willing to be matched to a subset of the objects of the matching at time step *t* + 1.
- the total number of objects at time step *t* might not equal the total object to which they are to be matched on the objects at time step *t* + 1.
- the resulting matching might not match all of the objects.

Since we do not have the equally sized sets of elements, each object may only be willing to be matched to a subset of the objects on the other side of the matching but a stable matching will still exist (i.e., matched results in the same preferential ordering from both sides, i.e., proposor and acceptor).

## III. Stable matching based on Voronoi partition

Here, we use a Voronoi diagram as a more robust and reliable metric to identify the mappable condition and thus enable track cells that undergo significant movement or exist in a densely populated region.

### A. Voronoi diagram

Given a set of *n* points (i.e., cell location, *p*_1_, *p*_2_, ⋯, *p*_*n*_ ∈ *P* in ℝ^2^), a Voronoi diagram partitions the 2-D plane into *n* Voronoi regions (i.e., *V*(*p*_*i*_) ≜ *V*_*i*_) with the following properties: 1) each point *p*_*i*_ lies in exactly one region and 2) if a point *q* ∉ *P* lies in the same region as *p*_*i*_, the Euclidean distances from *p*_*i*_ to *q* will be shorter than the Euclidian distance from ∀*p*_*j*_ ∈ *P* to *q*.

#### Definition 1

Given *n* points *p*_1_, *p*_2_, ⋯, *p*_*n*_ in a distance space (*S*, *d*), partition *S* into regions *V*_1_, *V*_2_, ⋯, *V*_*n*_ such that *V*_*i*_ = {*s* ∈ *S*|*d*(*s*, *p*_*i*_) < *d*(*s*, *p*_*j*_), *i* ≠ *j*. The resulting partition of space is called a *Voronoi diagram*.

In what follows, we will focus on Voronoi diagrams in Euclidean space so the distance metric will be *d*(*p*_*i*_, *p*_*j*_) ≜ ‖*p*_*i*_ − *p*_*j*_‖_2_. In general, a Voronoi region *V*_*i*_ is defined as the intersection of *n* − 1 half planes formed by taking the perpendicular bisector of the segment:

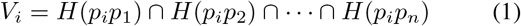

where *H*(*p*_*i*_*p*_*j*_) represent half plane formed by taking the perpendicular bisector of 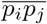.

A Voronoi cell encapsulates all neighboring objects that are nearest to each object and reflect two characteristics of a coordinate system: i) all possible points within one’s Voronoi cell are the nearest neighboring points for that object and ii) for any object, the nearest object is determined by the closest Voronoi cell edge.

### B. Object tracking using stable matching

In cell tracking, we have consecutive image frames *t* and *t*+1 time step where we have objects 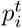, *i* = {1, ⋯, *n*} and 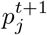, *j* = {1, ⋯, *m*} respectively. Proximity-based tracking algorithm sets parameter, i.e., cutoff length (*l*_cutoff_) so that one can determine whether two cells are mappable to each other by checking 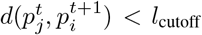. However, in general it is difficult to choose *l*_cutoff_. For instance, if the cell population is dense or the cell size is small, we need to choose small *l*_cutoff_. On the other hand, if the cell population is sparse or the cell size is large, we may need to choose large *l*_cutoff_ to link tracks.

#### Lemma 1

Consider consecutive frames *t* and *t* + 1 time steps and assume 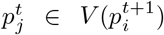. Then, 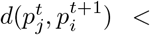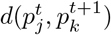 where superscripts denote time sequences and *k* ≠ *i*.

*Proof:* (by definition of Voronoi partition)

Followed by Lemma 1 and if there exists only one cell 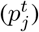 located in 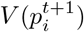, then 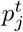 is more likely to be followed by the cell 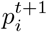 among many other cells 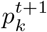.

Since a Voronoi diagram can subdivide the 2-D plane into polygons based on cell locations, there exist small polygons where the population is dense. On the other hand, where the cells are sparsely populated, there exist large polygons. Thus, it can automatically handle both sparse and dense population region and intrinsically, handle the size of the cell (i.e., as the cell size is bigger, the size of polygons is bigger since many cells cannot be packed in a small region).

Now we present how to use this to link tracks over time. In general, we can consider two scenarios. First, there exist at most one object 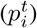 in 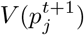 and then we can simply map 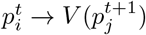 or 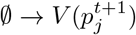. Second, we consider many-to-one matching case, i.e., there exist at least two objects 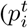, 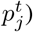 belong to the same Voronoi cell 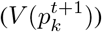. To address this, we consider a stable matching problem [29], [30].

Figure 1 shows an illustrative comparison of the proximity-based and Voronoi-based tracking approach in stable matching problem setting where we assume the radii of each ball (*B_i_*) satisfy 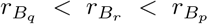. Preference relationship can be defined in terms of distance and then, from *a*, we have *d*(*a*, *q*) < *d*(*a*, *r*) < *d*(*a*, *p*) and from *b*, we have *d*(*b*, *r*) < *d*(*b*, *p*) < *d*(*b*, *q*). Although *b* prefers *r* to *p*, *r* prefers *a* to *b* based on the proximity-based preference relations, i.e., *d*(*a*, *r*) < *d*(*b*, *r*) < *d*(*b*, *p*). It is no difficult to prove this. Since *r* ∈ *B*_*p*_ but *r* ∉ *V*(*b*),

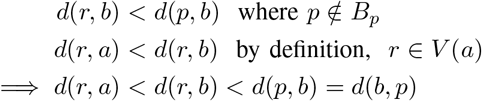

**Fig. 1.**
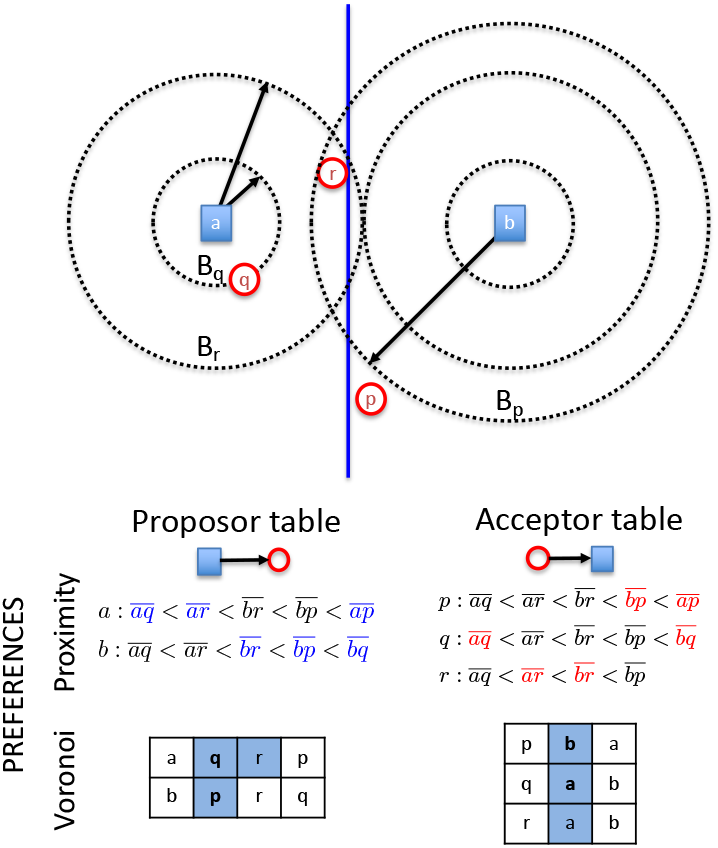
An illustration of proximity-based and Voronoi-based approach where (*a*, *b*) represents object at time step (*t* + 1) and (*p*, *q*, *r*) represents object at time step (*t*) and dot lines represent the boundary of bounding disk *B*_*q*_, *B*_*r*_ with center *a* and *B*_*p*_ with center *b* respectively. Solid blue line represents Voronoi edge between *V*(*a*) and *V*(*b*).

Thus, if we use proximity-based approach by setting cutoff length (*l*_*cutoff*_) such as *d*(*b*, *r*) < *l*_*cutoff*_ < *d*(*b*, *p*), we can find *l*′ such that *d*(*a*, *r*) < *l*′ < *d*(*b*, *r*) < *l*_*cutoff*_ < *d*(*b*, *p*) and thus, *r* should be assigned to *a* instead of *b* (i.e., not stable). To generalize this, we define a *blocking object* as follows:

#### Definition 2

Given a set of *n* points (*a*, *b*, ⋯) in ℝ^2^ and their Voronoi partitions (*V*(*a*), *V*(*b*), ⋯), a *blocking object* can be defined as follows: for a given *p* ∈ *V*(*b*), there exists *r* satisfying i) *r* ∈ *V*(*a*) and ii) preference metric *l*(*r* → *b*) < *l*(*p* → *b*) where *V*(*a*) and *V*(*b*) are adjacent to each other (i.e., sharing Voronoi edge) and *l*(*x* → *y*) represents a preference metric (*x*: proposor and *y*: acceptor).

As an example, for the proximity-based approach, preference metric can be defined as 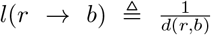, i.e., inversely proportional to the proximity distance. Then, based on proximity-based approach, blocking object does not allow stable matching followed by Definition 2, i.e., resulting in the different preferential ordering from both proposor and acceptor as shown in Figure 1.

#### Lemma 2

Given a point in ℝ^2^, if there exists any blocking object, proximity-based tracking (or nearest neighbor-based approch) cannot provide a stable matching.

*Proof:* The reader is referred to Figure 1 for a graphical description of this proof (simply ignore *q* here). As a proposor, *b* prefers *r* to *p* since *d*(*r*, *b*) < *d*(*p*, *b*) where *r* represents a blocking object and *p* represents nearest neighbor of *b* among any other objects in *V*(*b*). However, as an acceptor, *r* prefers *a* to *b* since *d*(*r*, *a*) < *d*(*r*, *b*) by definition of Voronoi partition (*r* ∈ *V*(*a*) where *V*(*a*) and *V*(*b*) are adjacent Voronoi partition), i.e., different preferential ordering from both sides (not stable).

For given objects set 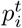 and 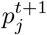 in frame *t* and *t* + 1, a Voronoi-based preference metric *l*(*x* → *y*) can be defined in terms of the conditional probability function. We denote *x* and *y* as objects from two consecutive time frame respectively. For instance, if *x* represents object from *t* + 1, then *y* represents object from *t* and vice versa. For a given *y*, the preference of *x* → *y* is defined as inverse of the probability of the joint of events *x* and *y* by measuring overlapped area between *V*(*x*) and *V*(*y*):

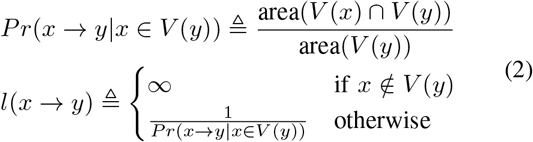

Here, for a given *V*(*y*), we measure the overlapped region between *V*(*x*) and *V*(*y*) as the metric of the conditional probability of *x* given *y*. Note that if *x* ∉ *V*(*y*), by definition *Pr*(*x* → *y*|*x* ∈ *V*(*y*)) = 0 and thus *l*(*x* → *y*) = ∞ which enables to remove blocking object in our consideration. If there exist more than one object in *x*_i_ ∈ *V*(*y*), preference metric is inversely proportional to the occupancy of Voronoi partition from the objects with respect to the overlap area, i.e., area(*V*(*x*_*i*_) ∩ *V*(*y*)).

We consider two scenarios of matching based on both proximity-based and Voronoi-based preference:

- Case 1 (equally sized set (bijection)-assuming that we do not have *q*): for the proximity-based approach, in the proposor table, we have 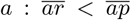 and 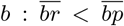 and in the acceptor table, we have 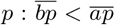 and 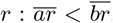. Thus, (*a*, *q*) results in the same preferential ordering but *b* prefers *r* to *p* and *r* prefers *a* to *b*, i.e., the first matched set prefers some given elements of the second matched set over the element to which is already matched (not stable match). On the other hand, based on Voronoi-based preference, there is no blocking object by definition (i.e., *r* ∈ *V*(*a*) should be assigned to *a* instead of *b* since *l*(*r* → *b*) = ∞). So, the matching results in the same preferential ordering from both sides, i.e., *a* ⇄ *r* and *b* ⇄ *p*.
- Case 2: different sized set (considering *q*) in this setting, since we do not have equally sized sets of elements, each object may only be willing to be matched to a subset of the objects on the other side of the matching but a stable matching will still exist (i.e., matched results in the same preferential ordering from both sides). Using the proximity-based approach, *r* is a blocking object so a matching is not stable. However, based on Voronoi-based approach, *a* can be matched to *q* although both *q* and *r* prefers *a* to *b* and (*b*, *q*) results in the same preferential ordering (i.e., matching is stable).

From these two observations, we illustrate that Voronoi-based preference enables stable matching although the proximity-based preference does not result in stable matching. To generalize this, we propose the following theorem:

#### Theorem 1

Based on Voronoi partition, a new preference metric (*l*) in equation (2) enables stable matching by removing blocking object.

*Proof:* For given adjacent Voronoi partitions *V*(*a*) and *V*(*b*), assume that there exists *r* ∈ *V*(*a*) such that *d*(*r*, *b*) < min_i_ *d*(*p*_*i*_, *b*) where *p*_*i*_ ∈ *V*(*b*). Since *r* ∈ *V*(*a*), *l*(*b* → *r*) = ∞ and *l*(*p*_*i*_ → *p*_*i*_) < *l*(*b* → *r*) ∀*p*_*i*_ ∈ *V*(*b*). Similarly, *l*(*p*_*i*_ → *b*) < *l*(*p*_*i*_ → *a*). Then, we have 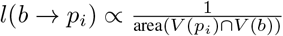 and 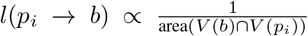, the preference ranking should be matched (thus stable matching *b* ⇄ *p*_*_ where *p*_*_ max_*i*_ area(*V*(*b*) ∩ *V*(*p*_*i*_)), *p*_*i*_ ∈ *V*(*b*)).

Followed by Theorem 1, we can guarantee stable matching in Voronoi-based tracking and we will describe a matching procedure in the following section.

### C. Algorithm

Consider time step *t* and *t* + 1 and assume that there exist *n* cells for time step *t* (i.e., 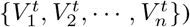) and *m* cells for time step *t* + 1 (i.e., 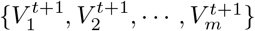) where 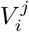 represents Voronoi partition corresponding to the *i*-th cell at time step *j*. Followed by Theorem 1, it simplifies tracking problem by breaking down into three cases: i) merging, ii) dividing and iii) one-to-one mapping as shown in Figure 2 and the each case can be described as follows:

- A) merging Voronoi partitions: due to the cell death or migration, more than one Voronoi partitions can be merged into one Voronoi partition (e.g., 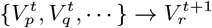) as shown in Figure 2A. We can simply match by using Voronoi-based preference metric or dominant region (i.e., 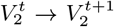 and 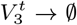) and then keep the previous label for tracking (i.e., 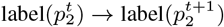) and 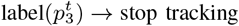).
- B) dividing Voronoi partitions: due to the cell division or migration (introducing a new cell), a Voronoi partition can be divided into more than two Voronoi partitions (e.g., 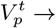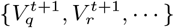). As shown in Figure 2B, we keep labeling for matched object based on Voronoi-based preference metric and for other object, we can annotate as a new label to keep tracking (i.e., 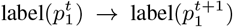) and 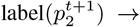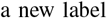).
- C) one-to-one map: as shown in Figure 2C, if we have one-to-one mapping 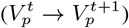, we simply track the previous label 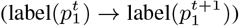.

**Fig. 2.**
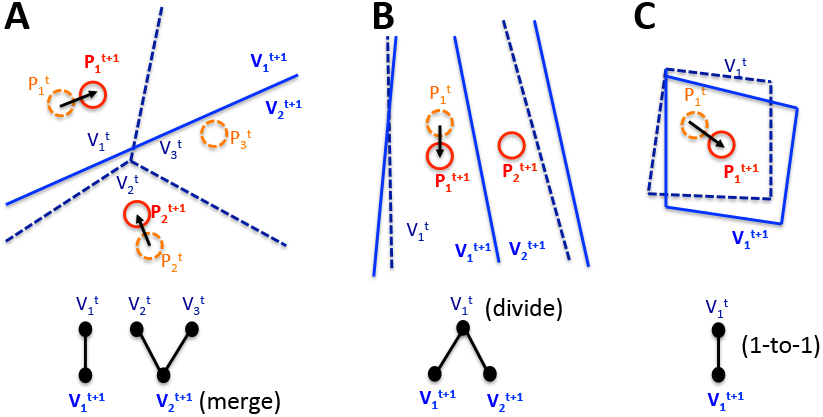
Conceptual illustration: A) merging, B) dividing and C) one-to-one mapping based on Voronoi partitions where solid red line indicates objects at time step *t*+1 and orange dash line indicates objects at time step *t*. Also, solid blue line and dash line represent corresponding Voronoi partitions at time step *t* + 1 and *t*.

To identify dominant region in A), we simply check the most frequent values (or area) in merged Voronoi (i.e., 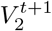) to map the Voronoi from the previous time step based on equation (2). For example, 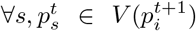 (i.e., *s* = {*j*, *k*, *l*, ⋯}),

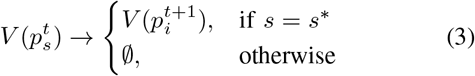

where 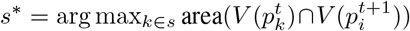. For the case in B), we first calculate 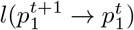 and 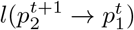 but since 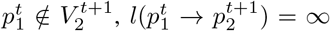. Thus, 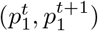 are matched as shown in Figure 2. Thus, we can simply check the condition whether each object belongs to Voronoi partition and select the corresponding Voronoi partition.

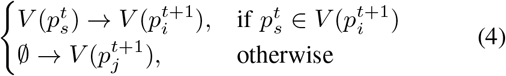

We summarize this procedure in **Algorithm 1**:

**Algorithm 1.**
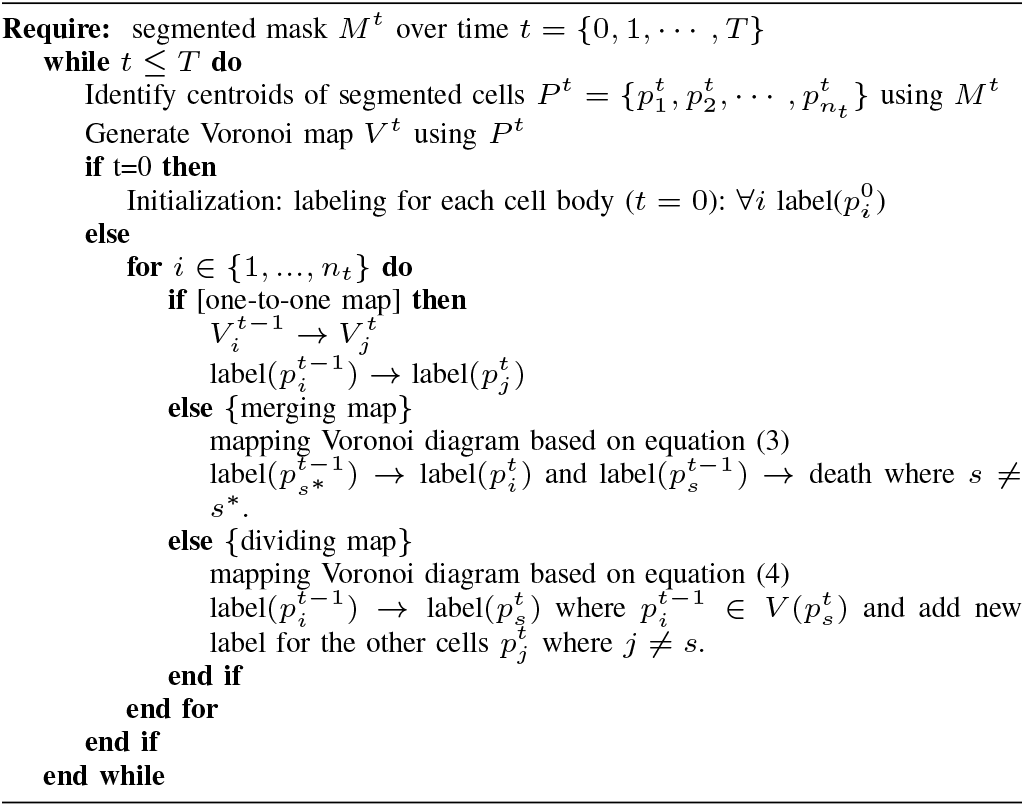
Voronoi-based object tracking

## IV. Results and Discussion

To evaluate performance, we simply generate synthetic trajectory in two consecutive time frame. Since we work in cartesian coordinates, for a given object, we generate the random angular and radial value (*ρ*) in polar coordinates and convert them into cartesian form. Since the tracking performance will be varying across different situations such as object density and speed (i.e., movement between consecutive time frame, *ρ*), we consider these parameters in simulation study. For example, Figure 3 shows examples of simulated trajectory: low density with small movement (left), low density with large movement (middle) and dense population with large movement (right). It is important to note that there exist a large overlap region of Voronoi partitions between consecutive frames, providing us more reliable and robust metric and improving tracking accuracy compared to proximity-based or nearest neighbor-based metric.

**Fig. 3.**
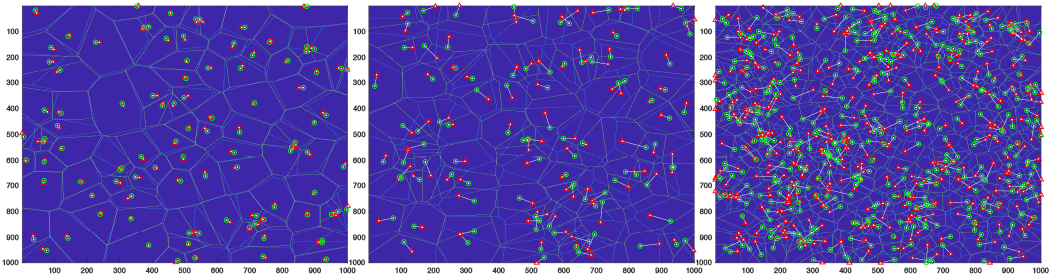
Examples of simulated trajectory for tracking task: (left) low density and slow movement (middle) low density and large movement (right) high density where green circle and red triangle represents objects from *t* = 0 and *t* = 1 frame respectively. Solid white line indicates true movement and blue and green polygon line indicate Voronoi partition of *t* = 0 and *t* = 1 respectively corresponding to green circle and red triangle object.

To compare tracking performance, we evaluate tracking accuracy with three different methods: i) proximity-based tracking, ii) nearest-neighbor based tracking and iii) Voronoibased tracking. Since we evaluate the performance with the numerical simulations as shown in Figure 3 (i.e., no object segmentation in our study), we do not consider overlap-based tracking here. Note that proximity-based tracking is essentially variation of overlap-based tracking (i.e., for a given object, selecting *l*_*cutoff*_ similar to the diameter of the object). We also evaluate the performance of proximity-based tracking with varying *l*_*cutoff*_. As *l*_*cutoff*_ increases, for a given object, we can find many objects and then we simply assign the nearest object (i.e., the same result as nearest neighbor tracking). For the nearest neighbor-based tracking, from a given object at time *t*, we simply assign the nearest object at time *t* + 1.

First, we compare tracking performance of three different methods with respect to different density. Figure 4 shows the tracking accuracy of three methods. For given *N* (density) and *ρ* (movement), we vary proximity radius (i.e., *l*_*cutoff*_) for proximity-based tracking. As an example, if we choose small *l*_*cutoff*_ relative to *ρ*, we do not expect the proximity based tracking achieve good tracking performance. On the other hand, if we choose *l*_*cutoff*_ large enough (i.e., close to *ρ*), we have better chance to improve tracking performance. Note that as the proximity radii increase, the tracking performance converges to the same as nearest neighbor-based tracking as we expected. Also, for the fixed cell movement parameter (*ρ*), as object density increases, the overall tracking performance degrades.

**Fig. 4.**
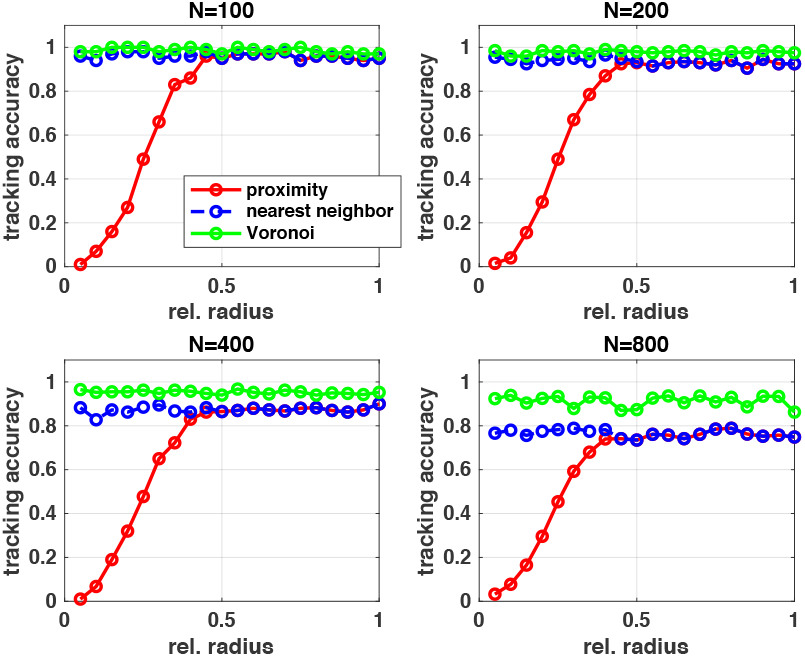
Tracking performance of three different methods: proximity-based, nearest neighbor-based and Voronoi-based approach across different object density scenarios where *N* represents the number of the objects in the same region. X-axis represents the relative radius (*l*_*offset*_/*ρ*).

Second, we compare tracking performance with varying cell movement. To do this, we simulate trajectories with different speed parameter (i.e., *ρ*) as shown in Figure 5 where we choose [*ρ* = *ρ*_*min*_ *ρ*_*max*_] for each object’s trajectory. As the movement increases, the tracking accuracy of nearest neighbor-based tracking approach decreases. On the other hand, the proposed approach shows better performance although the the overall performance decreases. Also, as the density increases (i.e., # of objects increases), the tracking accuracy decreases.

**Fig. 5.**
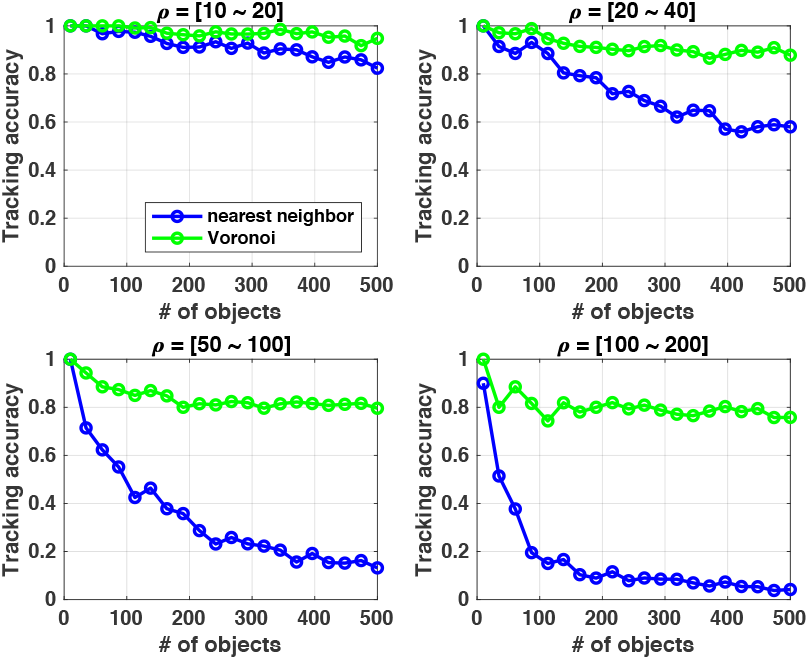
Tracking performance of nearest neighbor-based and Voronoi-based approach across different movement and density conditions.

It is important to note that Voronoi-based tracking only fails when there exists other object which has higher preference compared to the ground truth object since the proposed approach relies on only the position information. Therefore, if the cell migration speed is too high compared to temporal resolution, the proposed method will show the limited performance. To improve the tracking accuracy, we may need to consider complicating morphology-matching by using image features, i.e., similar cellular morphological features followed by the proposed method.

## V. Conclusion

We demonstrate Voronoi-based object tracking algorithm by considering stable matching problem shows better object tracking accuracy compared to proximity-based and nearest neighbor-based tracking. In the future work, we will apply the proposed method to neurodegenerative diseases application where we are manually curating longitudinal imaging data for evaluating tracking performance.

## Acknowledgements

This work was supported in part by the National Cancer Institute (U54CA209988, U2CCA233280).

